# Use of CD122+ natural killer cell precursors for superior anti-leukemia responses

**DOI:** 10.1101/2022.10.24.513585

**Authors:** Silvia Guglietta, Luis Cardenas, Carsten Krieg

**Author notes:** Corresponding author: Carsten Krieg, Dept. of Pathology,.

## Abstract

Acute Myeloid Leukemia (AML) is an aggressive blood cancer in adults. Intensive induction therapy successfully induces a complete response in up to 80% of the adult patients but a fraction of them is refractory or relapses. The fraction of non-responsive or relapsing patients is especially high amongst medically unfit and older patients, who cannot undergo aggressive chemotherapy treatment. Therefore, offering patients failing on first line intensive chemotherapy and medically unfit and older patients an effective treatment option for long-term control or even cure of disease represents a significant yet unmet clinical need.

As a component of the innate immunity, NK cells have recently shown great promise for the treatment of patients with different malignancies. Here, we show that injection of human interleukin-2 (IL-2) complexes (IL-2cx) induce the de novo generation and massive expansion of NK cell precursors *in vivo*. Furthermore, IL-2cx-expanded NK cells exert effector functions with the capacity to control the growth of non-self MHC class I-deficient RMA-S lymphoma cells *in vivo*, while remaining tolerant towards MHC class-expressing self RMA cells.

In an experimental setup mimicking the clinical case of refractory patients after intensive AML induction therapy, IL-2cx treatment mediated strong anti-AML responses following haploidentical bone marrow transplantation without the need for adoptive transfer of donor-derived or *in vitro* autologous expanded NK cells. Thus, this study demonstrates that IL-2cx immunotherapy allows the *in vivo* generation and expansion of functionally mature NK cells to achieve long-term response in AML paving the way to an effective treatment option.

## Introduction

Finding effective therapies against Acute Myeloid Leukemia (AML) has represented a daunting challenge, given the disease’s high fatality rate in adults^**1**^. A central issue in AML is that leukemia cells interfere with the production of normal blood cells in the bone marrow, causing weakness, infections, bleeding, and other complications ^2^.

AML is treated with remission induction therapy, an intensive form of chemotherapy, to reduce the leukemic burden and restore normal bone marrow function. Remission induction therapy depends on the patients’ medical fitness and is primarily determined by performance status and the nature and severity of medical comorbidities (e.g., heart disease, liver or kidney dysfunction, diabetes)^3^. Therefore, intensive chemotherapy for older patients (>75 years) is considered carefully.

The goal of care for medical-fit patients is to achieve long-term survival with the possibility of a cure. This requires complete AML remission. To achieve remission, for medically-fit patients with AML, the standard of care is the intensive “7+3” chemotherapy regimen. The 7+3 regimen is defined as 7 days of continuous infusion of cytarabine and 3 days of an anthracycline^4 5^. The intensive chemotherapy treatment generally causes three to five weeks of profound cytopenia and associated risk of lifethreatening infections and bleeding. In addition, many patients experience flu-like symptoms, nausea and vomiting, mucositis, alopecia, and diarrhea.

Within a medically fit population who can sustain the intense chemotherapy treatment about 60-70% of young patients will achieve a complete remission. This number is about 50% in adults older that 75 years. Yet, up to 20% of patients with a complete response will ultimately relapse and in the relapsing population the overall 3-year survival following salvage secondary chemotherapy is less than 10%^6,7^. Furthermore, this therapeutic regimen is a double hit to the patients because despite removing leukemia cells, it also removes antileukemia immune cells^8,9^. As a consequence, and in addition to losing anti-tumor efficacy due to therapy-induced immunosuppression, the patients become more susceptible to infections. Patients who undergo allogeneic hematopoietic cell therapy after showing a complete response with intensive chemotherapy appear to have a better outcome^6^. Allogeneic hematopoietic cell therapy is the greatest chance of attaining cure of patients with relapsed or refractory AML but also up to 20% of patients show remission^6,7^.

Several new approaches using Bite T cell therapy or anti-CD33 antibody therapy offered positive proof of concept that relapsing patients can be saved by novel therapeutic approaches and immunotherapy alone is sufficient for anti-tumor activity after overall response rates to these therapies were between 25% to 30%^10,11^. However, most patients do not respond, which highlights the urgent need to identify new therapeutic approaches.

Natural killer (NK) cells are cytotoxic innate lymphoid cells and are attractive targets for developing new cancer immunotherapies^12^. NK cells develop from bone marrow-derived progenitor cells in a thymus-independent manner^13^. By contrast with T and B cells, which express diverse antigen receptors generated by gene rearrangements mediated by recombination activating genes (RAG1/2), NK cells express a broad array of germline-encoded receptors^14,15^. IL-15 strongly stimulates NK cell proliferation, activation, and cytotoxicity and is essential for normal NK cell development and survival^16,17^. Adoptive transfer of purified NK cells has demonstrated efficacy in preclinical models of sarcoma, myeloma, carcinoma, lymphoma, and leukemias^18–27^. Indeed, NK cells mediate a graft-versus-leukemia effect against acute myeloid AML in patients receiving haploidentical bone marrow (BM)^28,29^. Donor-derived NK cells are licensed by donor MHC-I but not inhibited by host MHC-I. Moreover, unlike alloreactive T cells, such anti-tumor responses are usually not accompanied by notable graft-versus-host disease (GVHD).

Transplantation of haploidentical, T cell-depleted BM differing in a subset of MHC-I molecules and KIRs achieved the desired graft-versus-leukemia effect in patients^29,30^. In addition, co-transfer of interleukin-2 (IL-2)-activated mature NK cells from the same haploidentical, KIR ligand-incompatible donor showed superior anti-tumor immune responses. Interestingly, the provision of mature NK cells improves donor BM cells’ engraftment without aggressive conditioning regimens^28,29^. However, this approach has several shortcomings.

First, NK cells represent a small fraction of the peripheral white blood cells and donor leukapheresis is needed to achieve sufficient starting numbers for adoptive transfer. This has represented a major limitation for a widespread application of NK cell-based immunotherapies.

Second, NK cell preparations need T cell purge to avoid GVHD^30^.

Third, current protocols for selecting, expanding, and activating NK cells *in vitro* suffer from inter-individual variations in total NK cell counts and, most significantly, in anti-leukemia NK cell numbers^29,30^.

Lastly, upon adoptive transfer, NK cells are short-lived *in vivo* unless they receive signals from pro-survival cytokines, such as IL-2 and IL-15^24,30^. Thus, patients receiving NK cells need supportive treatment with low-dose IL-2 to maintain, expand and activate the infused cells^24^. Pilot studies of adoptive transfer of IL-2 activated NK cells demonstrated clinical response^31^. However, IL-2 immunotherapy may cause toxicity via immune cell stimulation and endothelial cell damage. Furthermore, while IL-2 can expand and stimulate natural killer (NK) cell and CD8 T cell antitumor responses, it also contributes to the expansion of regulatory T cells, which will dampen over time anti-tumor immune responses^32^.

Based on these premises, new strategies to exploit the potent and effective anti-tumor activity of NK cells without the above-mentioned drawbacks are needed to prolong the survival of patients experiencing relapse after allogeneic hematologic cell transplantation. IL-2 was the first common gamma-chain family cytokine to be discovered as a T-cell growth factor and later understood to function as an inducer of cytotoxic T-cells, NK cells and Treg cells^33^. IL-2 signals through three receptor components, IL-2Rα, IL-2Rβ, and IL-2Rγ (IL-2Rγ is also known as the common cytokine receptor γ chain, γc)^34^. These three proteins in different combinations form three classes of IL-2 receptors: The low-affinity IL-2 receptors consist solely of IL-2Rα (Kd ~ 10-8 M), the intermediate-affinity receptors contain IL-2Rβ + γc (Kd ~ 10-9 M), and the high-affinity receptors contain IL-2Rα + IL-2Rβ + γc (Kd ~ 10-11 M). The functional receptors are the intermediate- and high-affinity receptors; these forms of receptors both contain IL-2Rβ and γc, and heterodimerization of their cytoplasmic domains results in signaling. NK cells and CD8+ T cells constitutively express intermediate-affinity receptors, while the high affinity receptor is expressed on regularly T cells. Consequently, low levels of IL-2 are likely to induce an antiinflammatory response by expanding Tregs, while an immunostimulatory effect requires high-dose (HD) IL-2 and is associated with serious toxicities^35,36^. Nextgeneration IL-2 molecules for cancer therapy are being developed to harness the immunostimulatory effect of IL-2 while avoiding Treg stimulation and toxicity^37,38^.

In the present study we used human IL-2 complexes (IL-2cx) to target the intermediate-affinity IL-2 receptor for the *in vivo* expansion of donor NK cells following BM transplantation and evaluated the efficacy and feasibility of this immunotherapy approach for the treatment of AML. We found that complexing human IL-2 with anti-human IL-2 antibody (clone Mab602, IL-2cx) enhanced the *in vivo* bioactivity of IL-2. IL-2cx were efficient at amplifying mature NK cells from bone marrow precursors. *In vivo* expanded NK cells by using IL-2cx demonstrated strong cytotoxic activity and effectively avoided relapse in an aggressive AML mouse model.

## Results

### IL-2cx stimulates both precursor and mature NK cells

Although IL-15 strongly stimulates NK cell proliferation, activation, and cytotoxicity and is essential for normal NK cell development and survival^16,17^, IL-2 can substitute for IL-15 to support NK cell proliferation since the receptors for IL-2 and IL-15 share common β (CD122) and γ chains (CD132). Accordingly, IL-2 has long been known to directly stimulate the proliferation and activation of T cells and NK cells^39,40^. Originally IL-2 was used to expand white blood cells *in vitro*, so named lymphocyte-activated killer (LAK) cells^39–41^. Subsequent work found that LAK tumor destruction is mediated by NK cells^42^. However, the therapeutic use of IL-2 is limited due to its severe toxicity, which ranges from flu-like symptoms to severe capillary leak syndrome especially when used at high doses^43^. Moreover, IL-2 induces proliferation and activation of suppressive regulatory T cells^44^. To overcome these issues, various groups, including our own developed modified versions and formulations of IL-2 with improved toxicity profiles^44–48^.

Here, we directed human IL-2 to the CD122+ NK cells by complexing human IL-2 with the anti-human IL-2 monoclonal antibody MAB602^46^.

First, we evaluated the capacity of IL-2cx to expand NK cell precursors in different settings *in vivo*. In the BM of WT mice we found a low number of lineage (Lin)–CD122+ cells, out of which approximately 22% were NK1.1– DX5– precursor NK cells (pNK), while the majority were CD161+DX5 (CD49b+) mature NK cells (mNK). Upon administration of IL-2cx to WT mice, we observed a 15-fold increase in mNK cells and Lin– CD122+ BM cells (**Fig. 1A**). As previously determined, recombinant IL-2 was injected at an equal stimulatory concentration as in IL-2cx (termed high-dose IL-2, HD IL-2) or lower better tolerated dose (termed low-dose IL-2, LD IL-2)^46^. LD IL-2 led to a negligible increase of Lin– CD122+ BM cells, while HD IL-2 expanded Lin– CD122+ BM cells by about 3- to 4-fold (**Fig. 1A**). These data show that IL-2cx are 10 times more potent than HD IL-2 in expanding Lin– CD122+ BM cells in WT mice.

**Figure 1.**
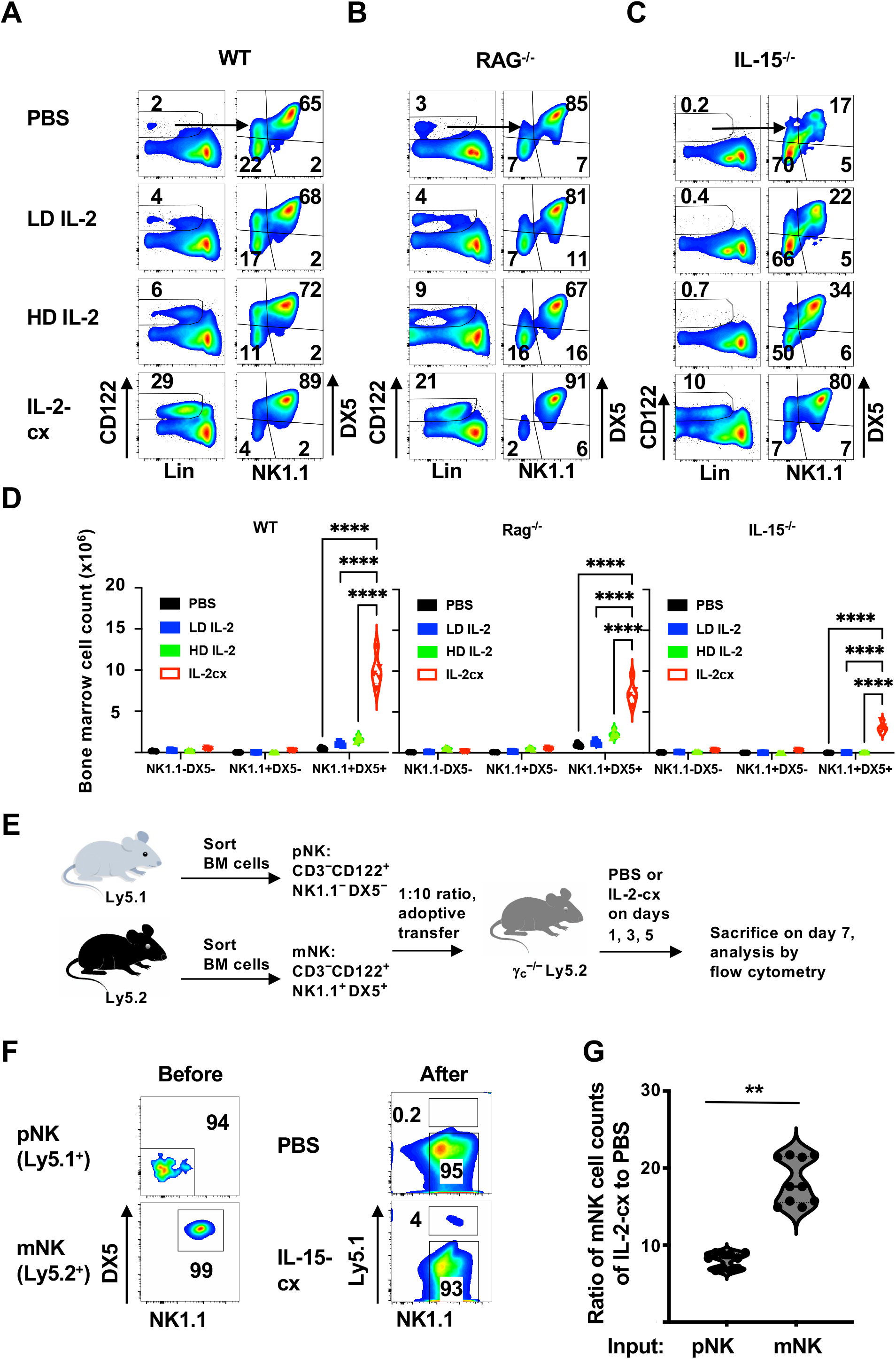
De novo generation of mature NK cells from precursors by IL-2 complexes. (**A**) WT, (**B**) RAG^−/−^, and (**C**) IL-15^−/−^ mice were treated with PBS, low-dose IL-2 (LD IL-2), high-dose IL-2 (HD IL-2) or IL-2 complexes (IL-2cx) for 5 days. DX5 and NK1.1 expression is shown in CD122^+^ lineage (CD3, CD4, CD8, CD19, Ter119, Gr1)-negative (Lin^−^) BM cells. (**D**) Columns represent mean cell numbers of indicated NK cell subsets. (**E**) CD3^−^ CD122^+^ NK1.1^−^ DX5^−^ precursor NK cells (pNK) from Ly5.1-congenic WT and CD3^−^ CD122^+^ NK1.1^+^ DX5^+^ mature NK cells (mNK) from Ly5.2-marked WT mice were sorted to a purity of 94% and 99%, respectively (**F**, left panels), followed by adoptive transfer at a 1:10 ratio to irradiated γc^−/−^ mice. Recipients were treated with PBS or IL-2cx on days 1, 3, and 5, and analyzed on day 7 for expression of Ly5.1 versus NK1.1 in CD3^−^ CD122^+^ splenocytes (**F**, right panels). (**G**) Ratios of donor-derived splenic mNK cells were calculated by dividing cell counts of donor derived mNK cells of IL-2cx-treated γc^−/−^ mice by those of PBS-treated γc^−/−^ animals. Data are representative of at least 2 independent experiments with a total of n=5-6 mice/condition. **P<0.01, ****P<0.0001.

The increase in Lin– CD122+ BM cells following IL-2cx treatment was not due to contamination by CD122+ T cells, as evidenced by a prominent population of pNK cells in RAG−/− mice (**Fig. 1B**). Consistent with published data, IL-15−/− animals showed near-normal counts of pNK but marked deficiency of mNK cells, these latter accounting for only 17% of Lin– CD122+ BM cells (**Fig. 1C**), therefore reflecting the dependency of mNK cell development and homeostasis on IL-15 signals^49^. Notably, in IL-15−/− mice treated with IL-2cx, the ratio of pNK to mNK cells became comparable to WT mice receiving IL-2cx, resulting in an 85-fold increase of mNK cell counts (**Fig. 1D**, bar graphs). Furthermore, similar to WT mice, administration of HD IL-2 but not LD IL-2 to RAG−/− and IL-15−/− animals caused noticeable expansion of Lin– CD122+ BM, which was 4- to 15-fold lower than with IL-2cx.

To obtain further evidence that IL-2cx stimulated pNK, we mixed highly-purified (94%) pNK from Ly5.1-congenic WT mice with 99% pure mNK cells from Ly5.2-congenic WT mice at a 1:10-ratio, followed by adoptive transfer to Ly5.2-marked common gamma chain (γc)-deficient recipients lacking mNK cells (**Fig. 1E**) ^50^. As shown in **Fig. 1E-F**, treatment of recipient γc−/− animals using IL-2cx led to an 8-fold expansion of donor mNK cells in the spleen compared to PBS-treated mice. This effect was even more pronounced for pNK cells, which expanded and efficiently differentiated into mNK cells upon IL-2cx, thus doubling the number of mNK compared to the PBS group.

In the periphery, upon injection of IL-2cx, NK cells underwent vigorous proliferation and expansion, peaking on day 2.5 in the blood and returning to near-normal levels on day eight after IL-2cx treatment (**Fig. 2A-B**). In addition to BM and blood, administration of IL-2cx also markedly expanded splenic NK cells in WT and RAG−/− animals (**Fig. 2C**). Furthermore, IL-2cx treatment led to a large population of NK1.1+ DX5+ mNK cells also in IL-15−/− mice (**Fig. 2C, right panel**), which generally lack mNK cells^49^. In contrast to IL-2cx, administration of recombinant IL-2 resulted in only minimal expansion of splenic mNK cells, even when IL-2 was given at high doses (**Fig. 2C**).

**Figure 2.**
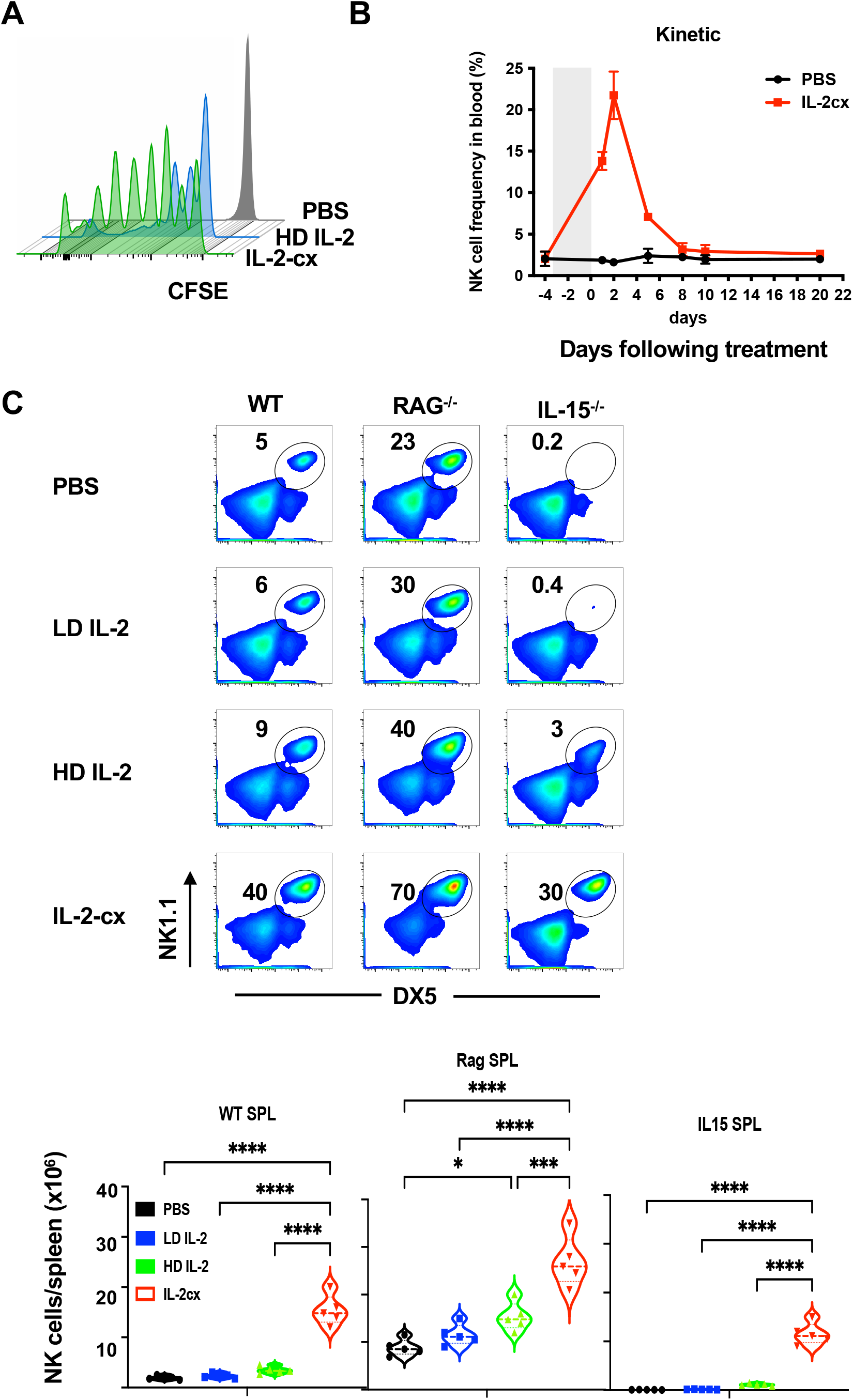
IL-2 complexes expand mNK cells. (**A**) Purified mNK cells from Ly5.1 - congenic WT mice were labeled using CFSE, followed by adoptive transfer to WT hosts receiving 3 daily injections of PBS or IL-2cx. Shown are CFSE profiles of Ly5.1^+^ donor mNK cells in spleen on day 4, with numbers indicating percentage of divided cells. (**B**) WT mice were treated with daily injections of PBS or IL-2cx for 3 days and frequencies of CD3^−^ NK1.1^+^ DX5^+^ mNK cells were analyzed in blood at indicated times. (**C**) Mice were treated as in **Fig. 1A**, followed by analysis of frequencies (top) and cell numbers (bottom) of NK1.1^+^ DX5^+^ mNK cells in spleen of WT, RAG^−/−^, and IL-2^−/−^ mice. Data are representative of at least 2 independent experiments with a total of n=5-8 mice/condition. n.s. not significant, *P<0.05, ****P<0.0001.

### IL-2cx activate NK cells

Besides supporting their proliferation, it is possible that IL-2 may also induce phenotypic and functional changes in NK cells. Therefore, we next examined the effect of IL-2 and IL-2cx on NK cell receptors and migration ability. As shown in **Fig. 3A**, compared to PBS and IL-2, treatment with IL-2cx resulted in increased levels of the inhibitory NK cell receptors Ly49G2 and NKG2A and the differentiation marker killer cell lectin-like receptor subfamily G member 1 (KLRG1) on mouse NK cells. These data suggest that upon IL-2cx treatment NK cells expand and become terminally differentiated.

**Figure 3.**
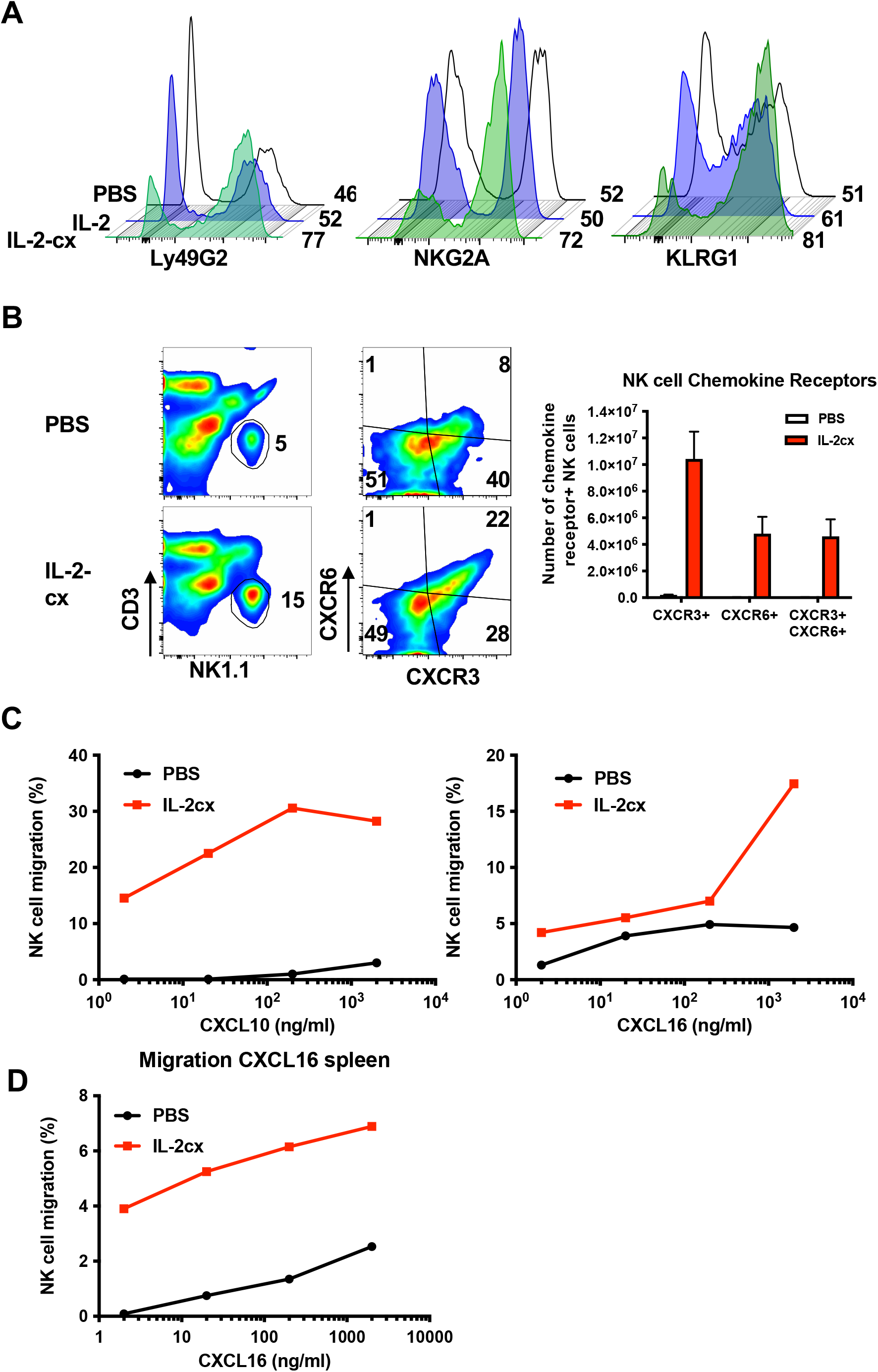
Characterization of mNK cells following IL-2 complexes. (**A**) WT mice receiving PBS, IL-2 or IL-2cx for 3 days were analyzed on day 4. Shown are profiles of Ly49G2, NKG2A, and KLRG1 in splenic CD3^−^NK1.1^+^ DX5^+^ mNK cells. (**B**, left) Flow cytometry analysis of chemokine receptor expression levels on WT and IL-2cx expanded mNK cells from spleen. (**B**, right) Total number of chemokine receptor positive cells per spleen. Migration experiments using a Boyden chamber: (**C**) Percentage of mNK cells isolated from spleen of WT or IL-2cx treated mice homing towards a CXCL10 or CXCL16 gradient and (**D**) percentage of mNK cells isolated from bone marrow homing towards CXCL16. Data are representative of 2 independent experiments with a total of n=2 mice/condition.

In addition to activating effector functions, an essential component of NK cell anti-tumor response is the ability to patrol tissues. To assess the homing capacities of *in vivo* IL-2cx expanded NK cells, we measured chemokine receptor expression on mNK cells (**Fig. 3B**). IL-2cx treatment up-regulated CXCR3 and CXCR6 chemokine receptors, which are both important for homing to the liver and bone marrow. Of note, especially bone marrow is a sites of leukemia cell residence in AML. To assess the functional significance of chemokine receptor up-regulation, we next performed an *in vitro* chemotaxis assay using Boyden chambers with purified mNK cells from the spleen of untreated or *in vivo* IL-2cx treated WT mice. As shown in **Fig. 3C**, we found that NK cells from BM of IL-2cx-treated mice more efficiently migrated toward CXCL10 (the ligand of CXCR3) and CXCL12 (the ligand for CXCR6). We obtained similar results using the mNK cells isolated from spleen (**Fig. 3D**).

### Increased NK cell effector functions following IL-2cx treatment

Our data showing that NK cells from IL-2cx-treated mice expressed a set of markers associated with activation led us to hypothesize that these cells would also be characterized by increased effector functions, including the ability to produce interferon-γ (IFN-γ) and exert cytotoxic activity^51^. In line with this prediction, highly purified splenic NK cells from IL-2cx-treated WT animals expressed higher granzyme B levels compared to IL-2 and PBS treated mice. Moreover, upon *in vitro* restimulation using titrated concentrations of stimulatory antibodies, NK cells from mice receiving IL-2cx produced significantly more IFN-γ than NK cells from PBS or IL-2-treated animals (**Fig. 4A, 4B**).

**Figure 4.**
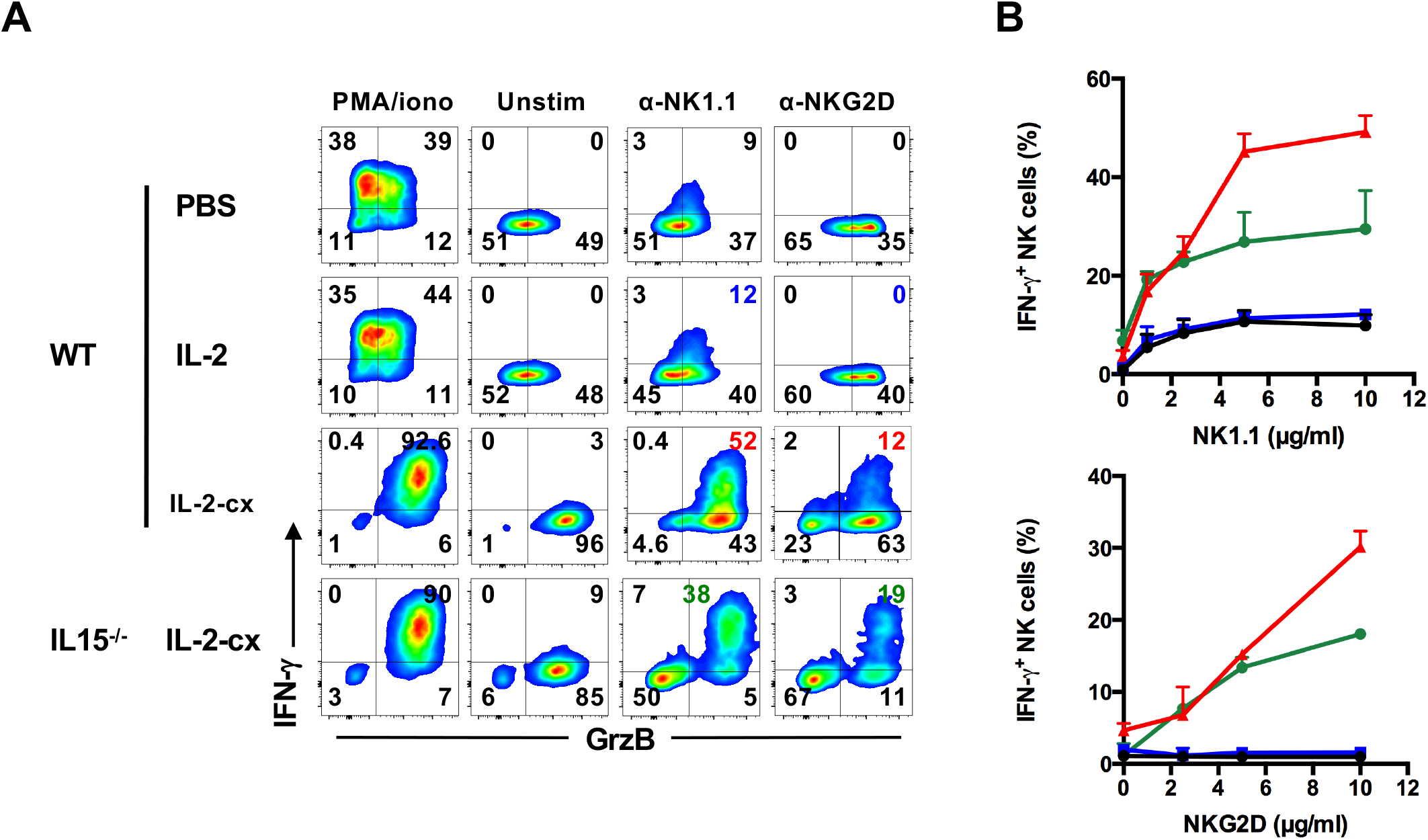
Production of effector molecules by IL-2 complex-activated NK cells. WT and IL-2^−/−^ mice were treated with PBS, IL-2 or IL-2cx for 3 days, followed by *in vitro* stimulation of highly-purified NK cells using plate-bound anti-NK1.1 (α-NK1.1) or anti-NKG2D (α-NKG2D) mAb for 5 hours, and intracellular staining for IFN-γ and granzyme B. PMA/ionomycin stimulation (PMA/iono) and unstimulated (Unstim) NK cells served as controls. Shown are dot plots of IFN-γ and granzyme B production of CD3− NK1.1+ DX5+ NK cells (**A**) and percentages of IFN-γ-secreting CD3^−^NK1.1^+^ DX5^+^ NK cells in response to titrated amounts of plate-bound α-NK1.1 or α-NKG2D (**B**). Data are representative of at least 2 independent experiments with n=5-6 mice/condition.

### IL-2cx-stimulates mNK cells to suppress tumor growth

Based on the data showing enhanced migration ability, higher activation measured by increased production of cytotoxic molecules and expression of inhibitory receptors in IL-2cx-stimulated NK cells, we next tested NK cell reactivity towards the syngeneic, MHC-I-proficient RMA T cell lymphoma cell line and the MHC-I-deficient RMA-S variant. To this aim, we subcutaneously injected mice with these cell lines and monitored tumor growth. WT mice transplanted with RMA cells were unable to control the tumor, regardless of whether they were treated with IL-2 or IL-2cx (**Fig. 5A**), thus demonstrating that NK cells remained tolerant to syngeneic cells upon IL-2cx treatment.

**Figure 5.**
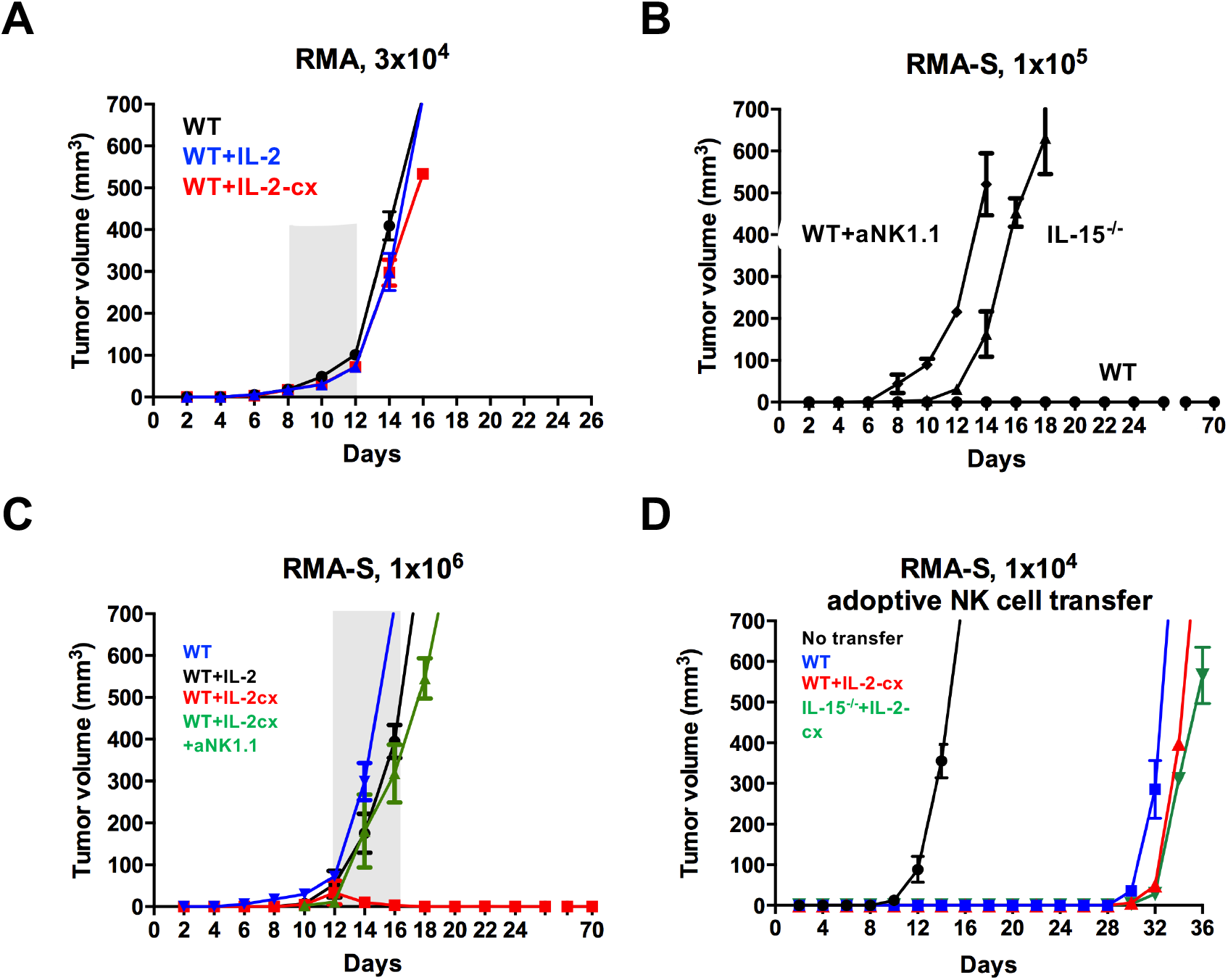
Efficient recognition of MHC-I-deficient tumor by IL-2 complex-stimulated NK cells. (**A**) 3×10^4^ MHC-I-proficient RMA cells were injected subcutaneously (s.c.) into WT mice and tumor volume in mm^3^ was measured every other day. When tumors were palpable, mice were treated with 5 consecutive injections of PBS (black), IL-2 (blue) or IL-2cx (red). (**B**) WT, NK cell-depleted WT (WT+α-NK1.1), and IL-2^−/−^ mice were injected s.c. with 1×10^5^ MHC-I-deficient RMA-S cells and tumor growth was monitored as in A. (**C**) 1×10^6^ RMA-S cells were injected s.c. into WT (black), WT receiving IL-2 (blue), WT receiving IL-2cx (red), and WT receiving IL-2cx plus α-NK1.1 mAb (green), followed by measurement of tumor volume as in A. (**D**) 1×10^6^ sorted NK cells from WT (blue) or WT (red) and IL-2^−/−^ mice (green) treated with IL-2cx for 3 days were adoptively transferred into irradiated (600 rad) γc^−/−^ recipients, followed by s.c. injection of 1×10^4^ RMA-S cells and measurement of tumor growth as in A. Data are representative of 2 independent experiments with n=6-9 mice/condition.

Conversely, MHC-I-deficient RMA-S cells were controlled in WT mice but multiplied in animals lacking mNK cells, such as IL-15^−/−^ and NK cell-depleted WT mice (**Fig. 5B**). When a 10-fold higher number of RMA-S cells was injected, tumor growth was evident in WT animals. Tumor control was restored upon IL-2cx treatment of WT mice. The anti-tumor effect depended on NK cells, as evidenced by a loss of tumor control when anti-NK1.1 mAb was given to deplete NK cells (**Fig. 5C**). Contrarily, IL-2cx treatment failed to induce long-term control of RMA-S in IL-15^−/−^ mice (data not shown), possibly indicating that tumor control was dependent on continued tumor suppression by mNK cells. The failure of IL-2cx-treated IL-15−/− animals to control RMA-S cells may result from decreased survival of the newly-generated mNK cells due to lack of IL-15. To assess this hypothesis, we generated mNK cells from WT, IL-2cx-treated WT, and IL-2cx treated IL-15^−/−^ mice and adoptively transferred them to γ_c_^−/−^ animals. γ_c_^−/−^ hosts contain normal to increased IL-15-levels^52^ and were transplanted subcutaneously with RMA-S cells. In contrast to controls, γ_c_^−/−^ mice receiving mNK cells suppressed RMA-S tumor growth for almost a month (**Fig. 5D**), suggesting that adoptively-transferred mNK cells reached replicative senescence at later time points or that tumor cells mutated, therefore escaping NK cell suppression.

### De novo generation of anti-leukemic NK cells by IL-2cx

To further assess the potential of de novo generated NK cells upon IL-2cx treatment, we conditioned C57BL/6 mice (H-2^b^) with either lethal (950 rad) or sublethal (600 rad) irradiation, followed by transplantation of NK and T cell-depleted BM from haplo-identical H-2^b/d^ F1(C57BL/6 x Balb/c) animals. While in mice receiving lethal irradiation 90% of splenocytes were of donor origin, sublethal irradiation of host mice resulted in less than 20% engraftment by donor cells (data not shown).

We next applied the above strategy to C57BL/6 mice that were sub-lethally irradiated and injected with a small number of C1498 AML cells (H-2^b^) to mimic residual disease following chemo- or radiotherapy. Simultaneously, these mice received either syngeneic (syn, H-2^b^) NK cell-depleted lineage-negative (ΔNK Lin^−^) or haplo-identical (haplo, H-2^b/d^) ΔNK Lin^−^ BM cells, followed by 3 cycles of IL-2cx. Control mice receiving only C1498 AML cells or sublethal irradiation died within 3 weeks (data not shown). Animals receiving sublethal irradiation along with ΔNK Lin^−^ haplo-BM cells plus PBS or HDIL-2 or mice getting sublethal irradiation plus ΔNK Lin^−^ syn-BM cells followed by IL-2cx showed a survival benefit of only 1-3 weeks (**Fig. 6**). Notably, mice receiving ΔNK Lin^−^ haplo-BM upon sublethal irradiation followed by 3 cycles of IL-2cx survived significantly longer, with about half of the animals surviving for more than 90 days (**Fig. 6**). This effect was dependent on CD122^+^ DX5^+^ NK cells, whereas CD8^+^ T cells were dispensable in this model (**Fig. 6**).

**Figure 6.**
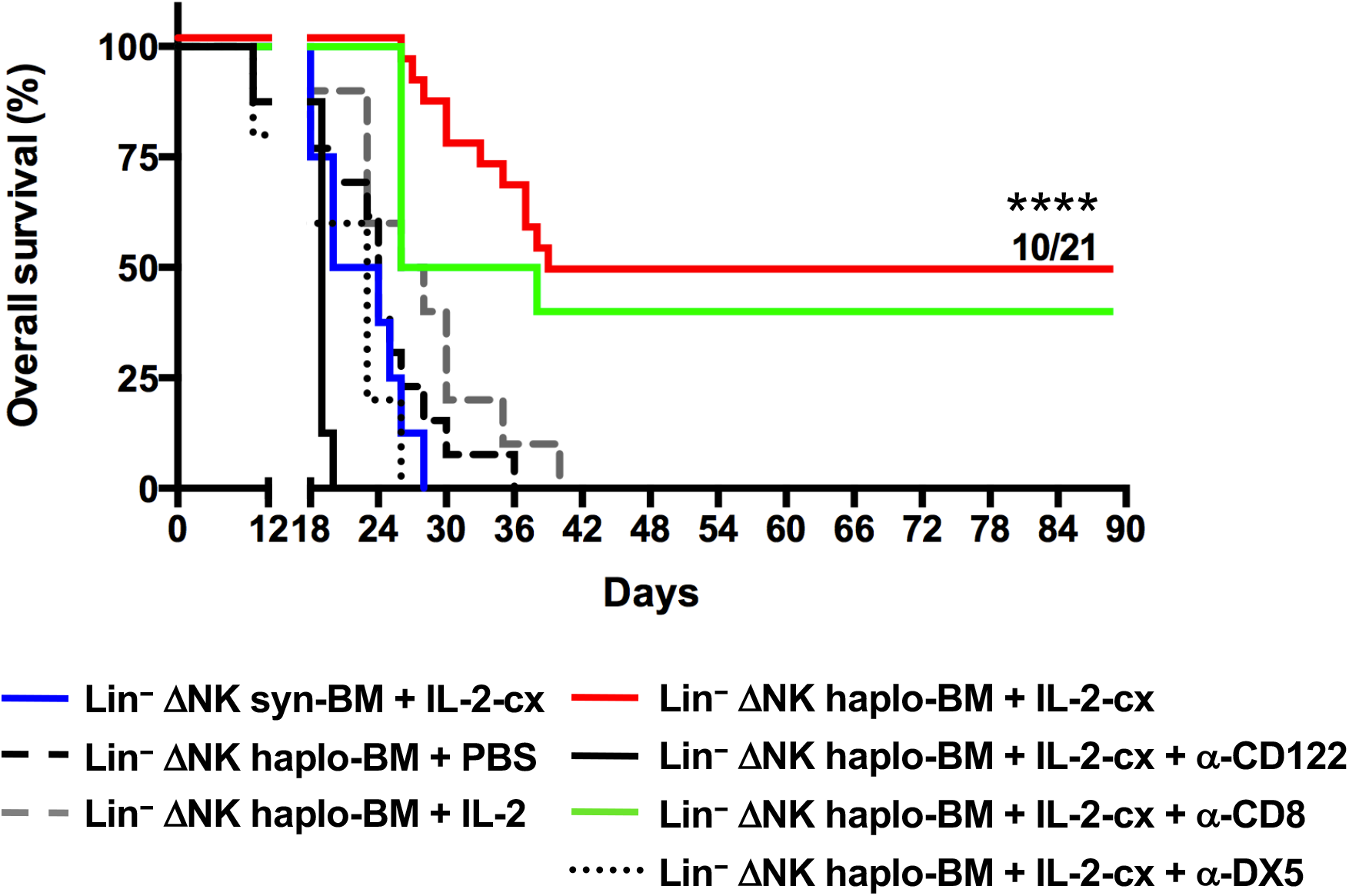
Anti-leukemic effects by newly-generated mNK cells from haplo-identical BM upon IL-2 complex treatment. Irradiated WT recipients (600 rad) were injected with 1×10^3^ C1498 myeloid leukemia cells along with 1×10^7^ NK cell-depleted Lin^−^ syngeneic (ΔNK Lin^−^ syn-BM, H-2K^b^) or ΔNK Lin^−^ haplo-identical BM cells (ΔNK Lin^−^ haplo-BM, H-2K^d/b^), followed by daily PBS, IL-2 or IL-2cx treatment on days 1-5, 8-13 and 16-20 after transplantation. Where indicated mice also received depleting anti-CD8 (α-CD8), anti-CD122 (α-CD122) or anti-DX5 (α-DX5) mAbs. In mice receiving ΔNK Lin^−^ haplo-BM plus IL-2cx, 10 out of 21 mice were still alive on day 90. n=5-21 mice/condition. ****P<0.001.

## Discussion

In this study, we report that a treatment course with IL-2cx can expand mNK cells and generate mNK cells from BM precursors. This finding is of interest for two reasons.

First, the ability of IL-2cx to drive NK cell development and maturation applied to both WT and IL-15^−/−^ mice, the latter of which usually lack mNK cells ^49,53^. Unlike endogenously-produced soluble IL-2, IL-2cx can compensate for the lack of IL-15 in facilitating normal NK cell development in IL-15^−/−^ animals. IL-2 activates CD122−γc receptor dimers by first binding to CD25, followed by CD122 and γc ^33^. IL-15 associates with the IL-15 receptor (IL-15Rα) already in cells producing IL-15, such as antigen-presenting cells (APC), followed by trans-presentation of IL-15–IL-15Ra complexes to CD122–γ_c_-expressing NK cells^54–57^. Notably, binding of IL-15 to IL-15Ra occurs with high affinity (dissociation constant Kd ≈10-11 M)^54–57^, which is necessary for IL-15 to be trans-presented, although non-immune cell-derived soluble IL-15Ra molecules can bind and neutralize IL-15^57,58^. This implies that the binding of CD122−γ_c_ dimers by IL-15−IL-15Ra complexes generates qualitatively similar responses in NK cells. This possibility is also supported by comparing the structures of the quaternary IL-2 and IL-15 complexes where no significant difference in CD122−γ_c_ receptor binding and signaling was reported between these cytokines ^59^. A recent clinical trial used the IL-15 agonist N-803 to support the expansion and survival of an autologous NK cell transplant one week after haploidentical BM transplantation in patients with AML^48^. Our work supports a simplified protocol meriting the treatment of patients with a CD122 agonist already upon haploidentical bone marrow transplant.

Second, the finding that IL-2cx generate new mNK cells from NK-committed BM precursors has important implications in the field of allogeneic hematopoietic transplantation. Previous research demonstrated that donor mNK cells facilitated donor BM engraftment. Thus, co-transfer of donor mNK and BM cells into sub-lethally irradiated hosts resulted in over 90% donor cell chimerism, similar to lethal irradiation of hosts; contrarily, sublethal irradiation alone was unable to achieve significant engraftment of donor cells^28^. Moreover, donor mNK cells protected against GVHD by killing host APCs that have been implicated in priming donor T cells to attack host cells^28^. Interestingly, as shown here, the use of IL-2cx obviated the need for co-transferring donor mNK cells to obtain >90% donor cell engraftment in sub-lethally irradiated haploidentical hosts. This result implicates that IL-2cx mediate rapid development and expansion of alloreactive donor NK cells, even in a lymphopenic environment. Moreover, this finding addresses a limitation of current *in vitro* protocols for obtaining NK cells. NK cell preparations vary considerably among patients both in total and leukemia-specific NK cell numbers^29,30^. Most importantly, our approach allows the patient specific tailoring of an NK cell therapeutic *in vivo* by using IL-2cx. When used in a murine model of AML, IL-2cx efficiently led to the development and expansion of donor BM-derived haploidentical NK cells, which mediated long-term survival of animals, while, unlike previous reports^60–62^, CD8+ T cells were dispensable in the herein used AML model.

Notably, treatment by IL-2cx did not appear to lead to loss of self-tolerance of NK cells as demonstrated by their ability to tolerate self-MHC-I-competent RMA cells while efficiently killing MHC-I-deficient RMA-S cells. The latter occurred very efficiently and might indicate reversal of an anergic state that NK cells adopt in the presence of MHC-I-deficient tumors, as recently shown^63^. Of note, as demonstrated by the increased expression of chemokine receptors, IL-2cx-generated NK cells showed improved migration abilities. Also, IL-2cx-stimulated NK cells did not exert cytotoxic activity against haemopoietic stem cells and tolerated normal hematopoietic cell lineages, as the counts of red blood cells, platelets, and various CD122– white blood cells remained unaffected upon IL-2cx administration.

Recent studies have demonstrated that NK cells can also control chronic myeloid leukemia^64^, and AML by *in vitro* purging and *in vivo* blockade of inhibitory NK cell receptors^65^. These data raise the possibility that IL-2cx treatment may be considered in hematologic malignancies as an adjuvant treatment with other NK cell-stimulating approaches.

## Materials and Methods

### Animals

C57BL/6 (WT), Ly5.1-congenic, RAG1^−/−^, IL-15^−/−^, γc^−/−^, D^b−/−^ K^b−/−^ (MHC-I^−/−^) mice (all on a C57BL/6 background), CB6F1 (F1 [C57BL/6 x Balb/c]) were purchased from Jackson. Mice were maintained under specific pathogen-free conditions at the animal facilities of the university hospitals and universities of Zurich and Lausanne. Experiments were performed following federal guidelines and under approved protocols at the universities of Zurich and Lausanne.

### Flow cytometry

Cell suspensions were stained using fluorochrome-conjugated mAbs targeting the following markers (from eBioscience and BD Biosciences unless otherwise stated): mB220 (RA3-6B2, Caltag), CD3 (145-2C11), CD4 (GK1.5), CD8 (53-6.7), CD11b (M1/70), CD11c (HL3), CD19 (1D3), CD25 (PC61), CD43 (1B11, BioLegend), CD49b (DX5), CD62L (MEL-14), CD69 (H1.2F3), CD122 (TMβ1), CD127 (A7R34), KLRG1 (2F1), Ly49D (4E5), Ly49G2 (4D11), Ly5.1 (A20), Ly6G (1A8), MHC-II (I-A/I-E, M5/114.15.2), NK1.1 (PK136), NKG2A (20d5), biotin-conjugated Ter119 (TER119) and biotin-conjugated Gr1 (RB6-8C5) along with phycoerythrin-conjugated streptavidin. Viable cells were acquired on a FACS Canto II, LSRII or LSR Fortessa (BD Biosciences) and analyzed using FlowJo software (TriStar Inc.). In some instances, mouse blood was analyzed using a differential blood picture analyzer (ADVIA, Siemens Healthcare).

### Cytokines and mAbs

IL-2cx, low-dose (LD) IL-2 (10’000 international units) and high-dose (HD) IL-2 (200’000 international units per injection) were prepared as previously described^46,66^, by complexing 1.5μg hIL-2 (National Cancer Institute) to anti-human IL-2 (15μg, clone MAB605) as indicated.

### Ex vivo degranulation and cytokine production

Murine NK cells were restimulated ex vivo with either phorbol myristate acetate plus ionomycin (PMA/iono, 50ng/ml and 1’000ng/ml, respectively), titrated amounts of plate-bound anti-NK1.1 (PK136) or anti-NKG2D mAb for 5 hours in the presence of brefeldin A (5μg/ml, Sigma). For intracellular cytokine staining of mouse cells, mAbs against IFN-γ (XMG1.2) and granzyme B (GB12, Caltag) were used, as previously established ^46^.

### Cell sorting and adoptive transfer of NK cells

Precursor NK cells (pNK) were purified by sorting CD122^+^ CD3^−^ CD4^−^ CD8^−^ CD19^−^ Gr1^−^ Ter119^−^ lineage (Lin)^−^ cells from Ly5.1-congenic BM cells. Where indicated NK cell-depleted Lin^−^ (ΔNK Lin^−^) cells were purified, consisting of DX5^−^ NK1.1^−^ Lin^−^ cells. Mature NK (mNK) cells were purified by sorting CD3^−^ CD122^+^ DX5^+^ NK1.1^+^ cells from spleen of Ly5.2-congenic mice using a BD Aria™ II cell sorter (BD Biosciences). At least 2×10^4^ purified pNK cells and/or 2×10^5^ purified mNK cells were injected intravenously into irradiated (600 rad) γ_c_^−/−^ hosts, as indicated. mNK cells from expanded WT or IL-15^−/−^ mice were sorted on day 2 after the last cytokine injection. In **Fig. 1F**, ratios were determined by dividing the counts of donor-derived mNK cells in spleens of animals receiving IL-2cx by those of mice given PBS.

### Tumor model in vivo

For the generation of subcutaneous tumors, 3×10^4^ MHC-I-competent RMA or 10^4^-10^6^ MHC-I-deficient RMA-S cells ^67^, both murine T cell lymphoma cell lines, were injected in 100μl RPMI into the upper dermis in the back of mice. Measurement of perpendicular diameters of tumors was performed every other day using a vernier caliper. Treatment was administered after tumors had reached a size of 25mm^2^ and became palpable. Where indicated mice received three weekly intraperitoneal injections of 100μg anti-NK1.1 mAb (PK136, BioXcell).

### Haplo-identical stem cell transplantation

H-2Kb-positive Ly5.1-congenic WT mice were conditioned using lethal (950 rad) or sublethal (600 rad) total body irradiation. The next day, 1×10^7^ T-cell-depleted BM cells with or without 1×10^6^ splenic NK cells from Ly5.2-congenic H-2Kb/d CB6F1 mice were injected intravenously, followed by treatment with PBS or IL-2cx as indicated.

### AML model

1×10 C1498 myeloid leukemia cells (ATCC) were mixed with 1×10 syngeneic or haplo-identical Lin– ΔNK BM cells and injected into sub-lethally (600 rad) irradiated B6 recipient mice followed by daily PBS, IL-2 or IL-2cx treatment on days 1-5, 8-13 and 16-20. Where indicated mice received 100μg depleting mAbs targeting CD122 (TMβ1), CD49b (DX5) or CD8 (YTS169).

### Statistical analysis

Data is presented following the ARRIVE guidelines^68^. Results were analyzed using GraphPad Prism 9.0 and are presented as mean ± standard deviation or standard error of the mean. Differences between groups were examined for statistical significance using one-way ANOVA with Bonferroni’s post-test or the student t-test. Survival curves were analyzed for significance using the log-rank test.

## Acknowledgments

Our work started in the Boyman lab in Switzerland, and we acknowledge feedback, especially from Onur Boyman, on the manuscript. In addition, we thank Werner Held for providing RMA cell lines and the animal facilities in Lausanne and Zurich for supporting our animal work. We are sponsored partly by the Cell Evaluation & Therapy Shared Resource, Hollings Cancer Center, Medical University of South Carolina (P30 CA138313), and an ACS-IRG to CK.

## Disclosures

The authors declare no conflict of interest related to this article.

## Authors Contribution

Authorship was decided following the COPE guidelines and based on substantial contributions which are listed as follows^69^: S.G. and C.K. conceptualized and designed the work. S.G. and C.K. acquired, analyzed and interpreted data. L.C. assisted in data analysis. S.G. and C.K. drafted the manuscript and revised it for intellectual content. All authors approved the final version of the article.

## Notes

### Competing Interest Statement

The authors have declared no competing interest.

### Summary of Updates

Updated Acknowledgements section.

